# β-endorphin primes Natural Killer cells and NK-derived Extracellular Vesicle to enhance anti-tumor cytotoxicity

**DOI:** 10.64898/2026.07.29.741417

**Authors:** Ofir Bar, Nofar Aharon, Muhammad Abu Ahmad, Ishai Luz, Olga Radinsky, Angel Porgador, Tomer Cooks

## Abstract

Psychoneuroimmunology suggests that positive physiological states, including laughter, could affect anti-tumor immunity, but the underlying mechanisms remain unclear. Here, we investigated whether β-endorphin (BE), an endogenous opioid peptide associated with positive physiological stimuli, modulates Natural Killer (NK) cell cytotoxicity and the anti-tumor activity of NK-derived extracellular vesicles (EVs). Using NK-92 cells, we assessed cytotoxicity against JIMT1 breast cancer cells, CD107a mobilization, cytotoxic activity of conditioned medium (CM), and EV yield, cargo, and function. BE enhanced NK-92-mediated killing of JIMT1 cells without increasing CD107a mobilization, suggesting that improved cytotoxicity was not driven by classical degranulation. Consistently, CM from BE-treated NK cells retained contact-independent cytotoxicity. NK-EVs were enriched in granzyme B and perforin following BE treatment exhibiting enhanced cytotoxicity against JIMT1 and BW tumor cells. BE also increased the cytotoxic activity of primary human NK cells, and BE-conditioned NK-EVs primed naïve NK-92 cells for enhanced tumor killing. These findings indicate that BE enhances NK anti-tumor immunity by remodeling the cytotoxic secretome and generating EVs that act as both direct cytotoxic effectors and mediators of NK cell priming.

**Graphical abstract:** 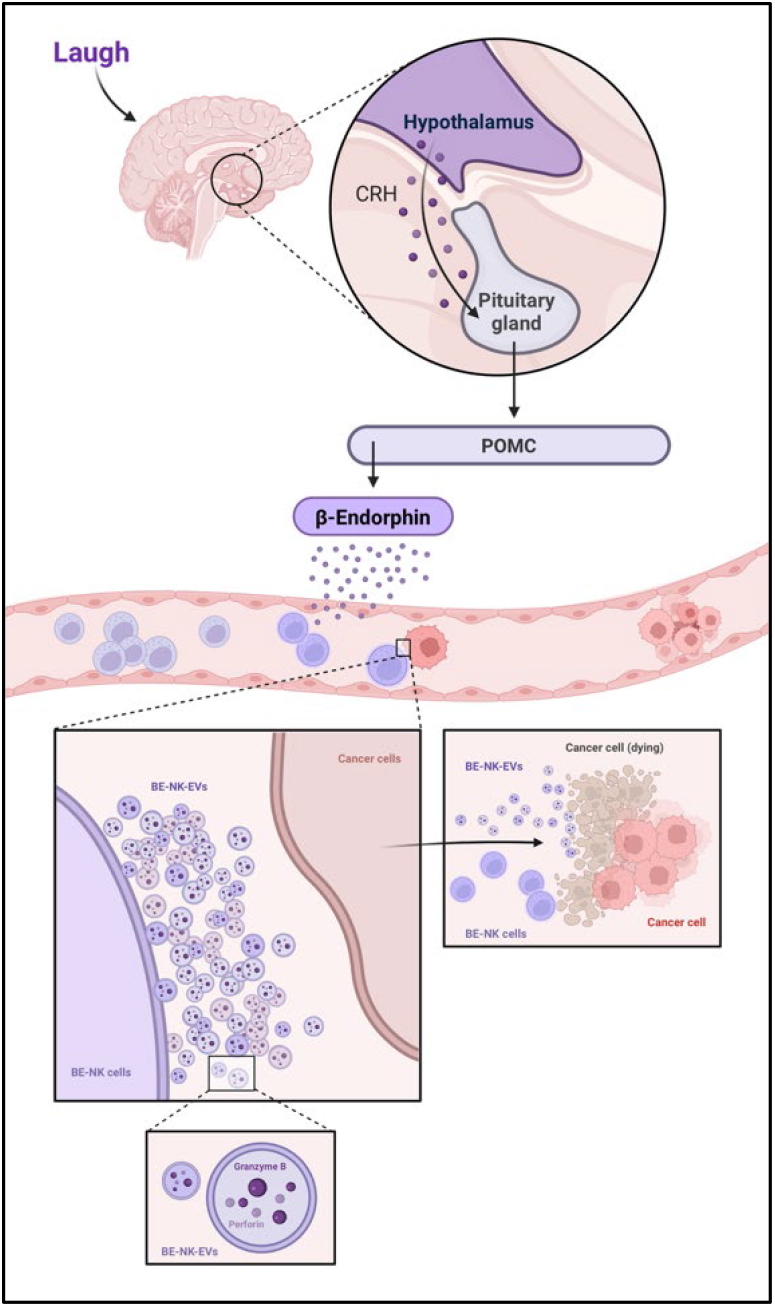

Proposed neuroendocrine–immune model linking positive physiological stimuli, NK cell-derived extracellular vesicles (EVs), and anti-tumor activity. Laughter is depicted as a conceptual upstream trigger of hypothalamic–pituitary signaling leading to β-endorphin (BE) release. BE conditioning enhanced NK-92 cytotoxicity and the anti-tumor activity of NK-derived EVs, consistent with granzyme B and perforin enrichment and supporting EV-mediated contact-independent cancer cell killing.

## Introduction

The interplay between psychological states and immune function, often termed psychoneuroimmunology, has long suggested that positive emotional experiences can translate into tangible physiological benefits.^1–5^ While the phrase “laughter is the best medicine” originated as an anecdote, accumulating evidence now supports its biological basis.^1,3^ Positive physiological stress, such as that induced by mirthful laughter, physical exertion, or deep relaxation, triggers a distinct neuroendocrine cascade that suggests active modulation of immune surveillance.^1,2,6–8^ Unlike chronic stress, which is often associated with immunosuppression, these positive states have been correlated with enhanced antiviral and anti-tumor immunity.^5,6,9^ However, while the systemic benefits of this “biology of hope” have been increasingly documented, the precise molecular mechanisms by which neuroendocrine signals translate into enhanced cellular lethality remain incompletely understood.

A key molecular candidate within this neuroimmune axis is β-endorphin (BE), an endogenous opioid peptide synthesized primarily in the pituitary gland and hypothalamus in response to various physiological stimuli, including stress, exercise, and positive affective states.^7,10–12^ While classically recognized for its analgesic and euphoric properties, emerging evidence suggests that BE may also function as a significant immunomodulator.^10–12^ This peptide is thought to bridge the nervous and immune systems by interacting with specific opioid receptors, most notably the mu (MOR) and delta (DOR) opioid receptors, which have been identified on the surface of various immune cell subsets.^7,13–16^ Among these, Natural Killer (NK) cells appear to be particularly responsive to opioid regulation.^7,8,14,16,17^ The presence of these receptors on NK cells suggests that they are primed to sense systemic neuroendocrine fluctuations, potentially translating signals from the central nervous system into alterations in immune surveillance and cytotoxic activity.^7,8,14–17^

NK cells are the cytotoxic lymphocytes that serve as guardians within the innate immune system, uniquely capable of recognizing and eliminating malignant or virally infected cells without prior antigen sensitization.^14,18–26^ Unlike T-cells, which require MHC restriction, NK cells are governed by a delicate balance of activating and inhibitory receptors, allowing them to rapidly attack “non-self” targets.^18–22^ Classically, NK cell cytotoxicity is mediated through the formation of an immunological synapse, followed by the polarization and exocytosis of lytic granules containing perforin and granzymes, which induce apoptosis in the target cell.^20,23,24^ Beyond direct killing, NK cells are potent producers of immunoregulatory cytokines (e.g., IFN-γ, TNF-α), thereby bridging innate and adaptive immunity.^20,21,25,26^ These intrinsic capabilities have positioned NK cells as a keystone of modern cancer immunotherapy, particularly in the development of “off-the-shelf” CAR-NK products for adoptive cell therapy.^19,20,22–25^ NK-92 cells provide a standardized and clinically relevant model for studying NK cytotoxicity and NK-derived products; however, validation in primary human NK cells is important to determine whether neuroendocrine modulation of NK function extends beyond an immortalized cell-line model.

While cellular therapies hold immense promise, their efficacy is often limited by the immunosuppressive tumor microenvironment (TME) and physical barriers that prevent immune cell infiltration.^25,27,28^ Consequently, attention has shifted toward the secretome, the complex array of bioactive molecules released by cells, as an alternative therapeutic modality. Central to this communication network are Extracellular Vesicles (EVs), membrane-bound nanoparticles (30–150 nm) constitutively released by almost all cell types.^25,26,29^ Far from being cellular debris, EVs serve as stable, long-distance messengers that transport a functional cargo of proteins, lipids, and nucleic acids, effectively mirroring the phenotype of their parent cell.^25,26,29^

EVs derived from NK cells (NK-EVs) have recently emerged as independent functional units with significant clinical potential.^25,26^ These vesicles inherit the cytotoxic machinery of the parent NK cell, enabling them to deliver “cytotoxic hits” to tumor cells without requiring direct cell-to-cell contact.^25,26^ As a therapeutic platform, NK-EVs offer distinct advantages over live cells: they are non-tumorigenic, have been reported to exhibit enhanced stability, can cross certain biological barriers (e.g., the blood-brain barrier), and may pose a lower risk of cytokine release syndrome.^25,26,29^

Despite the growing interest in NK-EVs as mediators of anti-tumor immunity, the upstream physiological signals that regulate their biogenesis and cargo loading remain largely unknown. Although it is well established that the immune system is influenced by the neuroendocrine system, the specific impact on NK cell vesicular output remains undefined. It remains an open question whether the neuroendocrine environment, specifically the surge of “laughter hormones” such as BE, can actively modulate the vesicular output of NK cells, potentially priming these vesicles for higher potency. Moreover, the precise cellular mechanisms by which BE modulates NK cell effector function remain understudied and are a primary focus of this investigation.

Here, we investigated how β-endorphin (BE) shapes the production and function of NK cell-derived extracellular vesicles (NK-EVs). Specifically, we examined whether neuroendocrine signaling remodels EV cargo and enhances EV-mediated anti-tumor activity while also assessing its effects on direct NK cytotoxicity. Finally, we validated the relevance of these findings in primary human NK cells. These studies aim to define how positive neuroendocrine signals enhance EV-mediated anti-tumor immunity and support the development of neuroendocrine-conditioned NK-EVs as a cell-free therapeutic platform.

## Methods

### Cell lines, primary NK cells and culture conditions

NK-92 cells were maintained at 37 °C, 5% CO₂ in α-MEM–based NK-92 medium supplemented with 12.5% heat-inactivated horse serum, 12.5% heat-inactivated fetal bovine serum (FBS), non-essential amino acids (NEAA), HEPES (N-2-hydroxyethylpiperazine-Nʹ-2-ethanesulfonic acid), 2 mM L-glutamine, 0.2 mM myo-inositol, 0.02 mM folic acid, 0.1 mM 2-mercaptoethanol, and penicillin–streptomycin. Recombinant human IL-2 was added to sustain NK-92 growth. Cells were kept in log-phase growth and split to maintain an appropriate density.

BW (hematologic) and JIMT1 (breast carcinoma) cell lines were maintained at 37 °C in a humidified atmosphere containing 5% CO₂. BW cells were cultured in RPMI-1640 medium, while JIMT1 cells were cultured in DMEM. Both media were supplemented with 10% heat-inactivated fetal bovine serum (FBS), 2 mM L-glutamine, sodium pyruvate, non-essential amino acids (NEAA), HEPES (N-2-hydroxyethylpiperazine-Nʹ-2-ethanesulfonic acid), and penicillin–streptomycin. JIMT1 cells were maintained as adherent monolayers and detached using Versene (EDTA) prior to flow cytometry-based assays, whereas BW cells were maintained in suspension. All cultures were routinely monitored for morphology and viability.

Primary human NK (pNK) cells were obtained from healthy donor peripheral blood and were enriched using RosetteSep according to the manufacturer’s instructions. Following enrichment, pNK cells were cultured in CellGenix SCGM medium supplemented with 10% human AB plasma and 300 IU/mL recombinant human interleukin-2 (IL-2). Cells were maintained at 37 °C in a humidified atmosphere containing 5% CO₂ until use in short-term cytotoxicity assays. Where applicable, human sample collection was performed under institutional approval and informed consent.

### Direct BE exposure during NK cell killing assays

To assess the effect of BE on NK-92 and pNK cells’ cytotoxic function under direct co-culture conditions, BE was added at the time of effector–target mixing (t=0) to achieve the indicated final concentrations (1 pM [10⁻¹² M] or 100 pM [10⁻¹⁰ M; 0.1 nM], depending on the experiment). UT co-cultures received an equal volume of the corresponding diluent and served as controls.

### Flow Cytometry-based cytotoxicity assay (direct co-culture)

Target cell killing by NK-92 and pNK cells was quantified using a flow cytometry-based viability assay. Target cells (BW and JIMT1) were harvested during log-phase growth, washed, and labeled with CFSE at a dilution of 1:1000 for 10 minutes to enable precise discrimination from effector cells during analysis. Labeled targets were then washed to remove excess dye and co-cultured with NK-92 effector cells in 96-well plates at effector-to-target (E:T) ratios of 1:1, 3:1, and 9:1. Co-cultures were incubated for 4 h and 18 h at 37 °C, 5% CO₂. At the endpoint, cells were stained with a viability dye, DAPI, at a concentration of 1:1000. For adherent JIMT1 cultures, targets were detached prior to acquisition. Flow cytometry data were acquired using Beckman coulter cytoFLEX S (model V4-B2-Y4-R3).

### Degranulation assay (CD107a surface mobilization)

NK-92 degranulation was assessed by measuring surface CD107a (LAMP-1)expression following exposure to BW or JIMT1 target cells. NK-92 effector cells were co-incubated with target cells at E:T ratios of 1:1, 3:1, and 9:1 in the presence of FITC-conjugated anti-human CD107a antibody (clone H4A3, mouse IgG1, κ; BioLegend, Cat# 328606). Co-cultures were incubated for 4 h or 18 h. Following incubation, cells were washed and analyzed by flow cytometry. NK-92 cells were gated according to forward and side scatter properties, and degranulation was expressed as the percentage of CD107a⁺ NK-92 cells.

### Conditioned medium (secretome) cytotoxicity assay

#### Preparation of NK-92 conditioned medium (CM)

NK-92 cells were seeded at a density of 1 × 10^6^ cells/mL in CM. Cells were treated with BE at final concentrations of 1 pM, 100 pM, 1 nM, or 1 µM, or left untreated (UT) as controls. Cultures were incubated for 24 hours at 37 °C in 5% CO₂ to allow accumulation of secreted factors. Following incubation, cell suspensions were centrifuged at 500 g for 5 min to pellet the cells. The resulting supernatants were collected and used immediately for functional assays.

#### Conditioned medium functional assay

JIMT1 target cells were harvested, washed, and seeded in 96-well plates at a density of 40,000 cells per well. To assess the dose-dependent effects of the secretome, defined volumes of NK-92 conditioned medium were added to the targets to achieve specific E:T ratios; for the 1:1 equivalent ratio, 40 µL of CM (derived from ∼40,000 NK cells) was added to the target wells. 3:1 equivalent ratio: 120 µL of CM (derived from ∼120,000 NK cells) was added to the target wells. Target cells were incubated with CM for 6 hours and 20 hours at 37 °C in 5% CO₂. At each endpoint, cells were harvested and stained with DAPI (1:1000).

#### ELISA

Soluble IFN-γ levels in CM were quantified using a custom sandwich ELISA. High-protein-binding 96-well plates were coated overnight at 4°C with an anti-human IFN-γ capture antibody (1:500 in Na₂HPO₄ buffer, pH 9.0). Plates were washed with PBST (PBS + 0.05% Tween-20) and blocked in PBST containing 10% FBS for 1 hour at 37°C. Following blocking, undiluted CM samples and a recombinant human IFN-γ standard curve (2,000-31.25 pg/mL; serially diluted in CM) were added to the plate and incubated for 2 hours at 37°C. Plates were washed, and a biotinylated anti-human IFN-γ detection antibody (1:500 in assay diluent) was added for 1 hour at 37°C. After additional washes, streptavidin-HRP (1:1,000) was added and incubated for 30 minutes. Signal was developed with the TMB substrate, and absorbance was measured at 650 nm with a microplate reader. IFN-γ concentrations (pg/mL) were calculated by interpolation from the standard curve generated in parallel.

#### BE conditioning for EV production

To generate BE-conditioned NK-92 EVs, human β-endorphin (β-endorphin 1–31; catalog no. RP11344; ≥95% purity, MW 3,465 Da) was purchased from GenScript (GenScript Biotech, Piscataway, NJ, USA). The lyophilized peptide was reconstituted in sterile distilled water to prepare a 10⁻^3^ M stock solution, aliquoted to avoid repeated freeze–thaw cycles, and stored at −20°C until use. Working solutions were freshly prepared by serial dilution in complete NK-92 culture medium immediately prior to each experiment.

NK-92 cells were cultured in EV-depleted conditions and treated with BE at final concentrations of 1 pM (10⁻¹² M), 1 nM (10⁻⁹ M), or 1 µM (10⁻⁶ M). Conditioned medium was collected for EV isolation as described below. EVs derived from untreated NK-92 cells processed in parallel served as controls (UT-NK-EVs).

#### EV isolation from NK-92 conditioned medium

EVs were isolated from NK-92 conditioned medium by differential centrifugation followed by ultracentrifugation. Briefly, supernatants were sequentially cleared to remove cells and debris: first, conditioned media were centrifuged at 1,500 RPM for 5 min at 4 °C to pellet intact cells, followed by a second centrifugation at 15,000 × g for 30 min at 4 °C (using rotor A50-8) to remove cell debris and larger vesicles. The clarified supernatants were then passed through a 0.22 µm vacuum filter (Millex-GP, Millipore, Cat# SLGP033RS). To isolate small EVs, the filtrate was subjected to ultracentrifugation at 38,700 RPM (∼150,000 × g) for 1.5 h at 4 °C using a Beckman Coulter OptimaTM ultracentrifuge with a 70 Ti rotor. The EV pellets were resuspended in sterile PBS (Gibco, Cat# 10010023) and subjected to a second round of ultracentrifugation under the same conditions (38,700 RPM, 1.5 h, 4 °C) to wash the vesicles and reduce soluble protein carryover. Final EV pellets were resuspended in sterile PBS and stored at −80 °C until use. EV preparations were derived from untreated NK-92 cultures (UT-NK-EVs) and BE-treated NK-92 cultures (BE-NK-EVs).

### EV quantification and characterization

#### Nanoparticle Tracking Analysis (NTA)

EV size distribution and particle concentration were measured by NTA (NanoSight NS500 instrument NTA 2.3) (Salisbury, UK). Samples were diluted in particle-free PBS to fall within the optimal detection range of the instrument. For each sample, multiple videos were acquired using consistent capture settings. Particle concentration (particles/mL) and modal size (nm) were exported for downstream normalization of EV doses in functional assays.

#### Western blotting

To compare the protein payload of EVs across treatment groups, samples were normalized based on particle count rather than total protein concentration. An equal number of EVs (as quantified by NTA) from each experimental condition was lysed with 5X sample buffer, boiled at 95 °C for 10 minutes, and loaded onto 12% self-cast Tris-Glycine SDS-PAGE gels. Proteins were separated by electrophoresis and transferred to nitrocellulose membranes (Greiner, Cat# 10-6000-02). Membranes were blocked for 1 hour at room temperature with 5% non-fat milk in PBS-T (0.1% Tween-20; Sigma-Aldrich, Cat# P1379) and then incubated overnight at 4°C with primary antibodies against EV-associated markers, including CD9 (polyclonal rabbit anti-human CD9; System Biosciences, Cat# EXOAB-CD9A-1; 1:1000) and ALIX (anti-human ALIX; [company], Cat# [catalog number]; [dilution]), as well as the cytotoxic effector proteins granzyme B and perforin. Granzyme B was detected using a mouse anti-human Granzyme B monoclonal antibody (clone #351927, Cat# MAB2906, R&D Systems/Bio-Techne) at 0.5 µg/mL (1:1000), and Perforin was detected using a mouse anti-human Perforin monoclonal antibody (clone #1031721, Cat# MAB103852, R&D Systems/Bio-Techne) at 2 µg/mL (1:250). Membranes were then incubated with the appropriate HRP-conjugated secondary antibodies, including HRP-conjugated donkey anti-mouse IgG secondary antibody (Cat# HAF018, R&D Systems/Bio-Techne, 1:1000), according to the host species of the primary antibody. Chemiluminescent detection was used to visualize protein bands. Densitometric quantification of granzyme B and perforin was normalized to the indicated EV-associated marker, CD9 or ALIX, and presented relative to untreated NK-EVs.

#### EV-Mediated cytotoxicity assay

EV-mediated killing was assessed by incubating target cells with particle-normalized EV doses quantified by NTA. UT-NK-EVs and BE-NK-EVs were added to BW and JIMT1 target cells at defined particle-to-cell ratios 10K:1, 30K:1, 90K:1, and 270K:1. Assays were performed in a fixed well volume (96-well, 160 µL/well) and incubated for 6 h and 20 h at 37 °C, 5% CO₂. At the endpoint, target cells were harvested, stained with a viability dye, and analyzed by flow cytometry to quantify target cell death. Target-only controls (no EVs) were included in each experiment.

#### Flow cytometry acquisition and analysis

Flow cytometry data were acquired on a calibrated cytometer. Target cells were labeled with carboxyfluorescein succinimidyl ester (CFSE) using the CellTrace™ CFSE Cell Proliferation Kit (Invitrogen, Thermo Fisher Scientific, Cat# C34554) prior to co-culture. Samples were acquired by flow cytometry, identified by forward/side scatter characteristics, and gated on CFSE⁺ target cells, and the percentage of dead targets (viability dye⁺, DAPI at 1:1000). Specific lysis was calculated as: Specific lysis (%) = 100 × [(% dead in co-culture − % dead in target-only) / (100 − % dead in target-only)]. Target-only wells (spontaneous death) and effector-only wells were included as controls in each experiment. For CD107a assays, NK-92 cells were gated based on scatter characteristics, and CD107a positivity was quantified using fluorescence-minus-one (FMO) controls where applicable.

#### BE-NK-EV priming of naïve NK-92 cells

To assess whether BE-conditioned NK-92-derived EVs can modulate the functional state of naïve NK-92 cells, a secondary EV-priming assay was performed. Naïve NK-92 recipient cells were harvested during log-phase growth, counted, and seeded in 6-well plates at 2 × 10⁶ cells per well in a final volume of 2 mL EV-depleted NK-92 medium supplemented with IL-2, corresponding to a final density of 1 × 10⁶ cells/mL.

Recipient NK-92 cells were incubated for 24 h at 37°C and 5% CO₂ with either PBS vehicle, β-endorphin (BE) alone, EVs derived from untreated NK-92 cells (UT-NK-EVs), or EVs derived from NK-92 cells conditioned with BE at 1 pM (10⁻¹² M), 1 nM (10⁻⁹ M), or 1 µM (10⁻⁶ M). For the BE-only control, BE was added to achieve a final concentration of 1 µM. EV treatments were normalized according to particle concentration measured by NTA, and all EV-treated wells received an equal number of particles. In the pilot experiment, EVs were added at 2.5 × 10¹⁰ particles per well, corresponding to approximately 1.25 × 10⁴ particles per recipient NK-92 cell.

Following the 24 h priming period, NK-92 cells from each condition were collected separately, transferred to tubes, and washed twice with PBS to remove unbound or residual EVs. Cells were then resuspended in fresh complete NK-92 medium, counted, and viability was assessed. The post-wash viable cell count was used to normalize effector cell input in downstream functional assays. To evaluate whether EV priming altered NK-92 cytotoxic capacity, washed recipient NK-92 cells were co-cultured with CFSE-labeled target cells at the indicated effector-to-target ratios. Target cell death was quantified by viability dye staining and flow cytometry, as described above.

### Statistical analysis

Statistical analyses were performed using GraphPad Prism. For experiments comparing ≥3 groups, one-way or two-way ANOVA with appropriate multiple-comparisons correction was applied as indicated in the figure legends. Data are presented as mean ± SD. The number of independent biological replicates (n) and technical replicates per experiment are reported in the figure legends. A p-value < 0.05 was considered statistically significant. All experiments included a minimum of three biological repeats unless stated otherwise.

## Results

### BE enhances NK-92 cytotoxicity without increasing CD107a degranulation

To determine whether physiologically relevant BE signaling can directly enhance NK cell cytotoxic function, we first evaluated the impact of BE conditioning on NK-92–mediated cytotoxicity against breast cancer cells. The NK-92 cell line, in particular, has emerged as a clinically relevant platform due to its unlimited proliferative capacity and high cytotoxicity against a broad spectrum of malignancies. To this end, NK-92 cells were treated with physiological concentrations of BE (1 pM and 100 pM) and co-cultured with JIMT1 breast cancer cells.

At the 4-hour time point, we observed a significant increase in tumor cell death in the BE-treated groups compared to untreated controls (Figure 1A). We used either a 1:1 or a 3:1 cell-type ratio (NK to JIMT1) and observed a highly significant enhancement at both effector-to-target ratios (p < 0.0001). We then validated this observation in two additional independent biological replicates (Figure 1B). This summary analysis confirmed that BE treatment consistently boosts NK cytotoxicity, resulting in a significant increase in killing capacity relative to basal levels. Notably, this immunostimulatory effect of BE was sustained over prolonged co-culture. In an 18-hour assay (Figure 1C, 1D), BE-treated NK cells maintained superior cytotoxicity, particularly at the 100 pM concentration. Notably, the 100 pM condition consistently drove the highest fold-increase in specific lysis across the dataset (p < 0.0001), indicating a dose-dependent reinforcement of anti-tumor activity over time.

**Figure 1.**
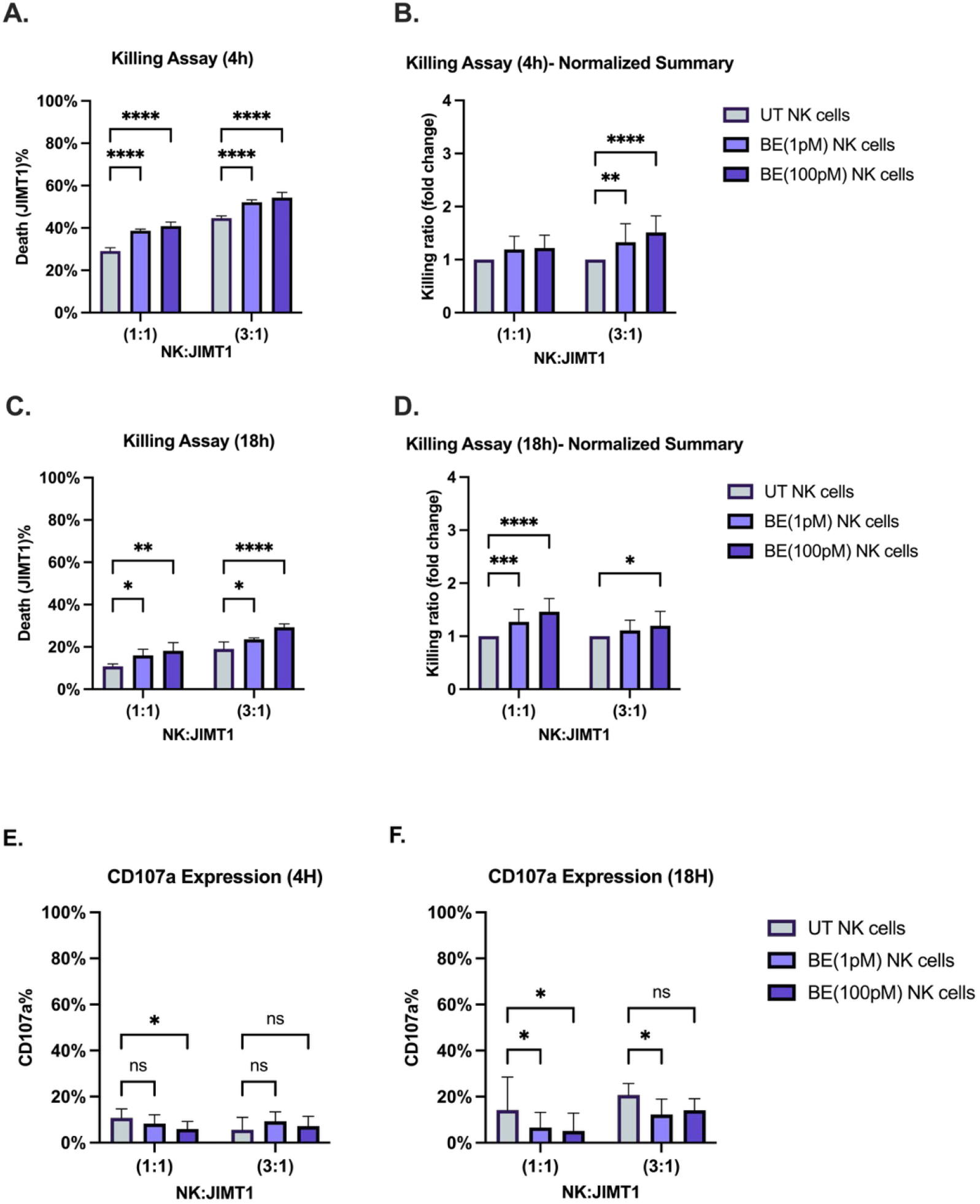
BE enhances NK-92-mediated cytotoxicity without increasing CD107a degranulation. NK-92 cells were pre-treated with BE (1 pM or 100 pM) or left untreatedof. NK-92 were then co-cultured with JIMT1 target cells. (A) Representative experiment showing the percentage of dead tumor cells after 4 hours of co-culture. The data represent 4 repeats. (B) Summary of cytotoxicity at 4 hours, presented as killing ratio (fold change) normalized to untreated NK-92 control. The data represent 12 repeats. (C) Representative experiment showing the percentage of dead tumor cells after 18 hours of co-culture. The data represent 4 repeats. (D) Summary of cytotoxicity at 18 hours, presented as killing ratio (fold change) normalized to untreated NK-92 control. The data represent 12 repeats. The percentage of CD107a⁺ NK-92 cells after 4 hours (E) and 18 hours (F) of co-culture. The data represent 16 repeats. Statistical analysis was performed as indicated in the Methods.

To determine whether the enhanced anti-tumor activity of BE-primed NK cells is driven by increased granule exocytosis, we assessed surface mobilization of the lysosomal marker CD107a (LAMP-1) in NK-92 cells following co-culture with JIMT1 targets. Unexpectedly, despite the increased killing capacity observed in cytotoxicity assays, BE treatment did not enhance CD107a surface expression. At the 4-hour time point, treatment with 1 pM BE resulted in a significant reduction in degranulation compared to untreated controls, particularly at the 1:1 effector-to-target ratio (Figure 1E). A similar trend was observed after 18 hours of co-culture, where BE-treated NK-92 cells again showed reduced CD107a expression relative to untreated controls, most notably at the 1:1 ratio (Figure 1F). Together, these findings suggest that the enhanced killing activity induced by BE is not accompanied by increased classical degranulation and may instead involve an alternative cytotoxic mechanism, potentially through secretion-associated pathways.

To determine whether the increased target-cell death observed in BE-treated co-cultures could be explained by a direct effect of BE on tumor cells, JIMT1 cells were exposed to the same BE concentrations in the absence of NK cells. Direct BE exposure did not induce a comparable increase in JIMT1 cell death relative to untreated controls (Figure S1). These data indicate that the enhanced cytotoxicity observed in NK-92 co-cultures is unlikely to result from direct BE-mediated toxicity toward JIMT1 cells, and instead supports an NK cell-dependent effect.

### BE-conditioned NK-92 secretome mediates contact-independent cytotoxicity independent of IFN-γ secretion

To assess whether the enhanced anti-tumor activity of BE-primed NK cells is mediated primarily by secreted factors rather than direct cell-to-cell contact, we evaluated the cytotoxic potential of cell-free CM. JIMT1 target cells were incubated with CM derived from NK-92 cells treated with a broad range of BE concentrations (1 pM, 100 pM, 1 nM, and 1 µM) (Figure 2A, 2B). CM derived from NK-92 cells was sufficient to induce tumor cell death, and CM from BE-treated NK-92 cells was associated with greater cytotoxicity than CM from untreated controls, consistent with the enhancement observed in direct co-culture assays. At 6 hours, this effect showed a dose-related increase in cytotoxicity, and at 20 hours, the cytotoxic activity was maintained and further increased. Together, these findings support the interpretation that BE promotes a contact-independent cytotoxic program within the NK-92 secretome.

**Figure 2.**
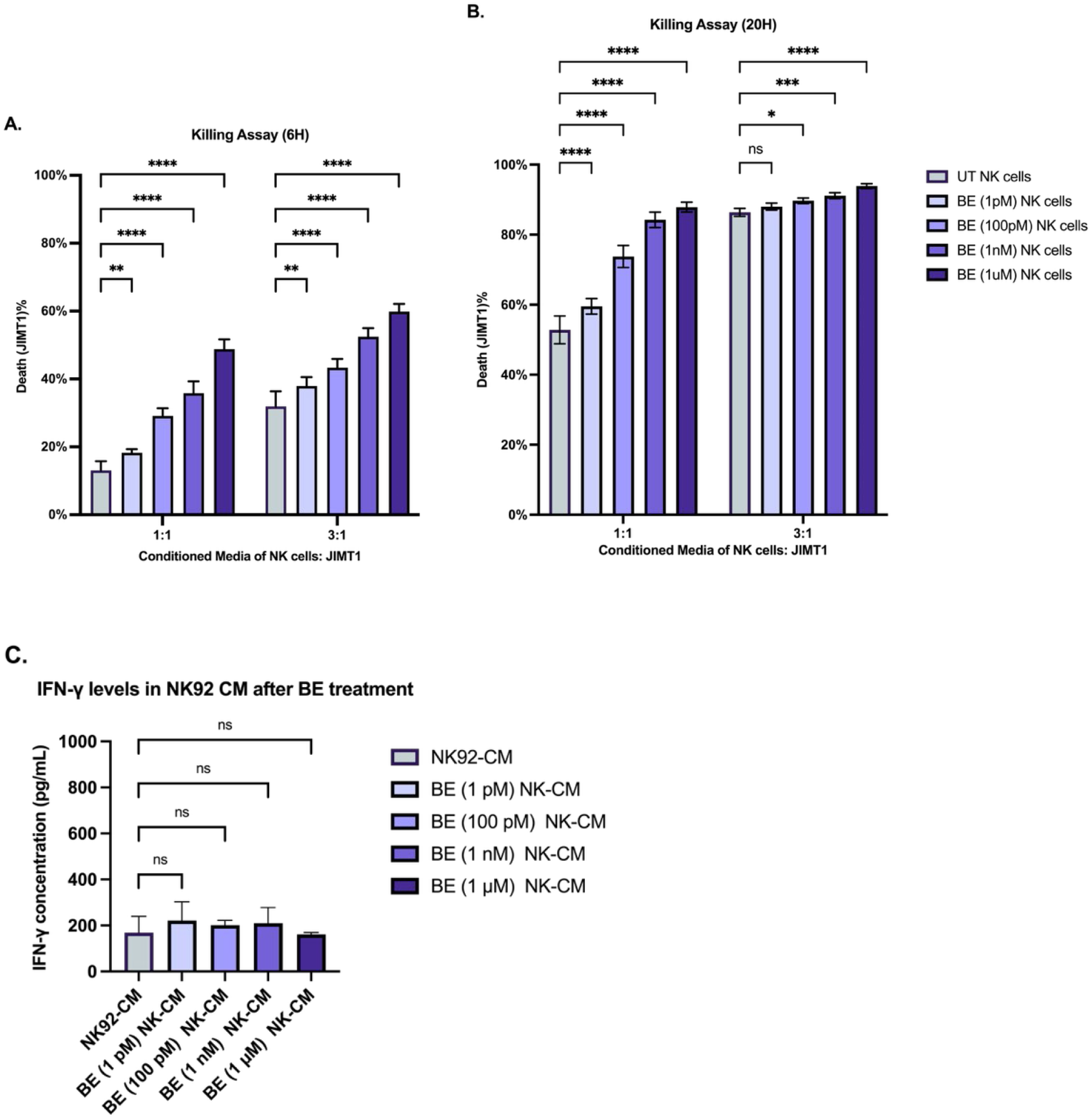
BE-conditioned NK-92 secretome mediates contact-independent cytotoxicity without detectable increase in IFN-γ. (A) Percentage of dead JIMT1 cells after 6 hours of incubation with CM collected from untreated or BE-treated NK-92 cells. The data represent 6 repeats. (B) Percentage of dead JIMT1 cells after 20 hours of incubation with the indicated CM. The data represent 6 repeats. (C) IFN-γ concentration (pg/mL) measured by ELISA in CM collected from NK-92 cells following 24 hours of incubation with the indicated BE concentrations or untreated control. The data represent 3 repeats.

To examine whether the increased anti-tumor activity observed in NK cells stimulated by BE secretome could be associated with a broad increase in cytokine release, we quantified IFN-γ levels in the CM using ELISA (Figure 2C). IFN-γ is a prototypical NK cell cytokine that can contribute to anti-tumor effects, including anti-proliferative signaling in tumor cells.^30,31^ Across the tested BE concentrations, IFN-γ levels in the CM were comparable to those measured in untreated NK-92 CM, and no statistically significant differences were detected between conditions. While basal IFN-γ was readily measurable, BE treatment did not show a clear trend toward increased secretion. Together, these data suggest that CM-mediated cytotoxicity is unlikely to be explained by changes in bulk soluble IFN-γ alone and are consistent with the possibility that other components of the secretome, including vesicle-associated factors, contribute to the observed functional enhancement.

### BE drives cytotoxic cargo loading without altering EV Biogenesis

Given our findings that BE, enhanced tumor killing persisted even in the absence of direct NK, target cell contact, we next sought to determine whether this effect is mediated by secreted factors, particularly EVs, as potential non-canonical cytotoxic effectors. To address whether BE influences NK cytotoxicity through changes in EV output or EV composition, we generated EVs from untreated and BE-conditioned NK-92 cells and characterized them by NTA and western blotting (Figure 3A).

**Figure 3.**
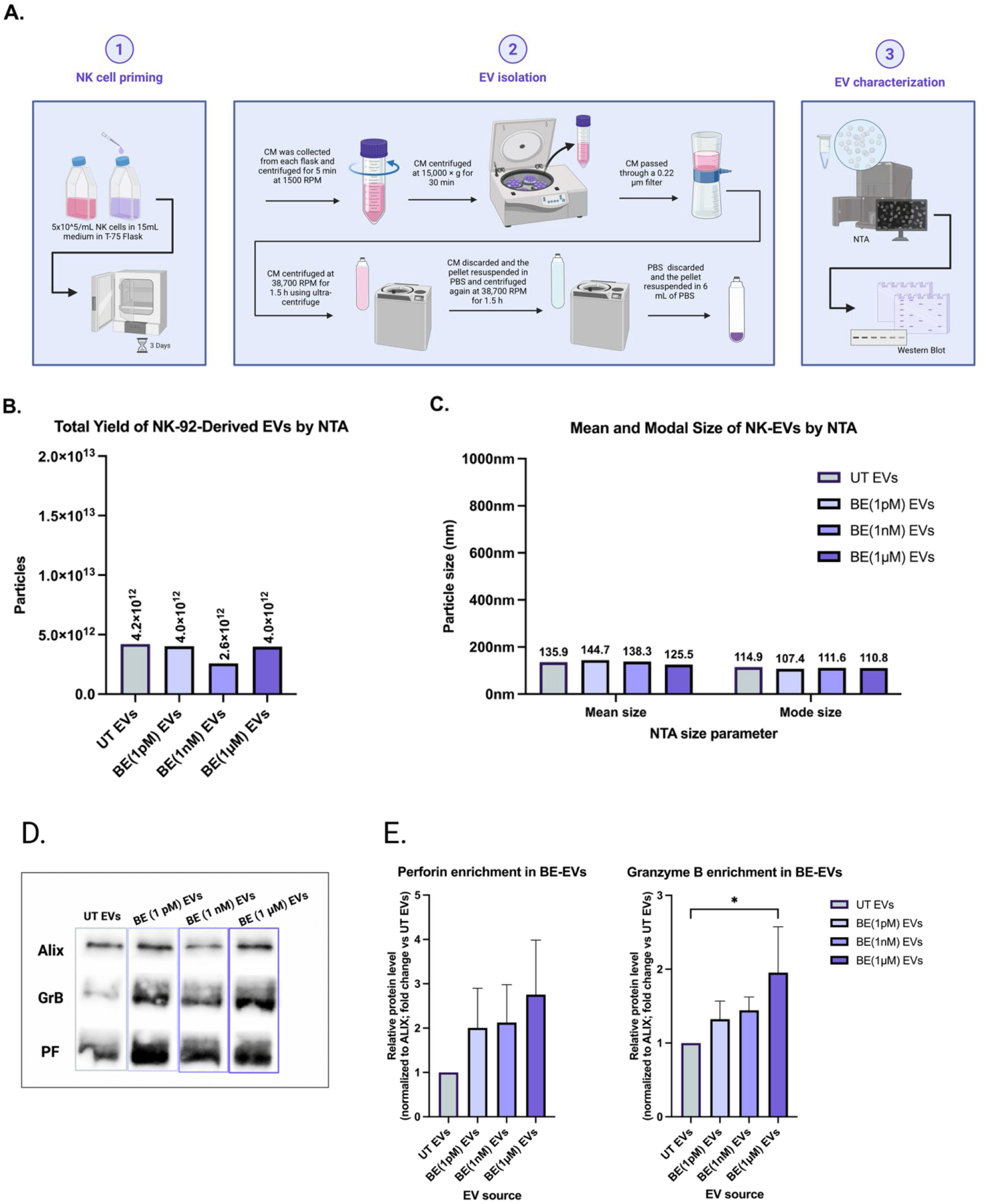
BE alters EV cargo composition but not total biogenesis. (A) Schematic overview of the experimental workflow for BE-EV generation, isolation, and characterization. NK-92 cells were primed with BE, conditioned medium was collected, and EVs were isolated by sequential centrifugation, filtration, ultracentrifugation, washing, and resuspension in PBS. Isolated EVs were then characterized by nanoparticle tracking analysis (NTA) and western blot. (B) NTA quantification of total EVs isolated from NK-92 cells following conditioning with escalating doses of BE (1 pM, 1 nM, 1 µM) or untreated control (UT). The bar graph displays the total particle count obtained per standard harvest. (C) Mean and modal particle size of EV preparations measured by NTA. (D) Representative western blot analysis of particle-normalized EV preparations isolated from UT and BE-treated NK-92 cells. Membranes were probed for the EV-associated marker ALIX and the cytotoxic effector proteins granzyme B (GrB) and perforin (PF). (E) Densitometric quantification of perforin and granzyme B levels normalized to ALIX and presented as fold change relative to UT EVs. Granzyme B was significantly enriched in EVs derived from the 1 µM BE condition, whereas perforin showed an increasing trend.

First, we analyzed the concentration of EVs isolated from untreated and BE-conditioned NK-92 cells using NTA (Figure 3B). The analysis confirmed that NK-92 cells maintain robust vesicle production regardless of BE exposure. The total EV yield remained comparable across the full BE concentration gradient, with untreated controls and cells treated with 1 pM, 1 nM, or 1 µM BE producing similar particle numbers per harvest. We next examined EV size distribution. Both the mean and modal sizes of NK-derived EVs were similar across all treatment groups (Figure 3C). These findings indicate that BE conditioning does not substantially affect the overall physical size profile of the isolated EV populations.

To investigate the molecular basis for the enhanced potency of BE-EVs, given that total particle yield remained constant, we characterized the protein cargo of the released vesicles using western blot analysis, focusing on the main cytotoxic molecules associated with NK cells: granzyme B (GrB) and perforin (PRF) (Figure 3D). ALIX detection was used as an EV-associated normalization marker. Western blot analysis of particle-normalized EV preparations revealed increased cytotoxic cargo following BE conditioning. Quantification of band intensities normalized to ALIX showed that granzyme B levels were elevated in BE-EVs, reaching statistical significance in EVs derived from the 1 µM BE condition (Figure 3E). Perforin showed a similar dose-associated increasing trend. This accumulation followed a clear dose-dependent trajectory: while 1 nM treatment induced a noticeable increase in effector cargo, EVs derived from the high-dose 1 µM condition displayed the highest abundance of both GrB and PRF. These data indicate that while physiological stress does not alter the quantity of EVs released, increasing neuroendocrine stimulation drives the preferential packaging of cytotoxic machinery into the EVs.

### BE conditioning potentiates EV cytotoxicity against solid and hematologic targets

To determine whether the BE-associated enrichment of cytotoxic cargo translates into enhanced functional activity, we performed particle-normalized EV killing assays against two distinct tumor models: JIMT1 and BW (hematologic malignancy) (Figure 4).

**Figure 4.**
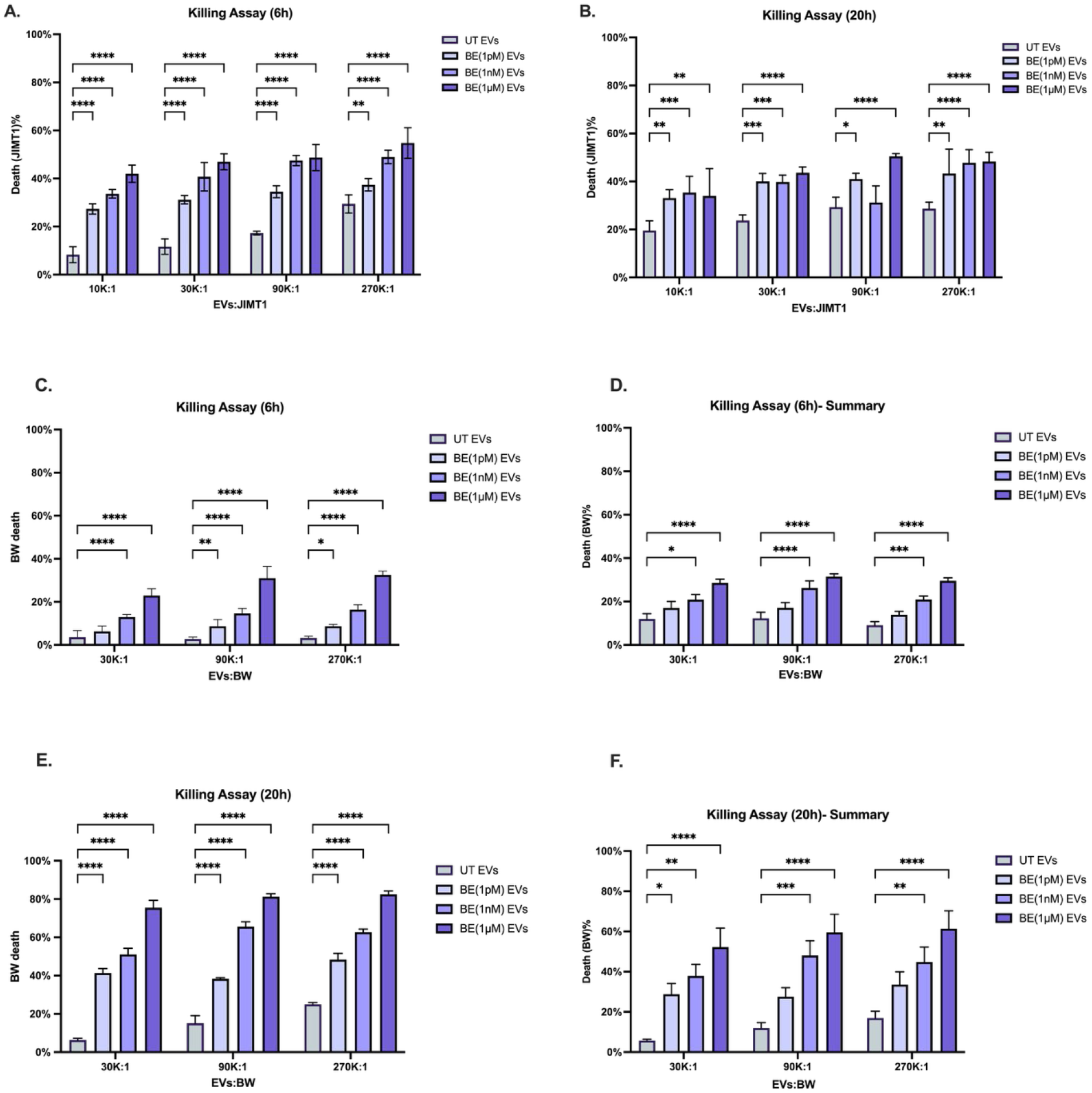
BE conditioning enhances NK-92 EV-mediated cytotoxicity. Particle-normalized EVs were isolated from untreated NK-92 cells (UT) or cells conditioned with increasing doses of BE (1 pM, 1 nM, 1 µM). EVs were then co-cultured with target cells at the indicated vesicle-to-cell ratios. (A, B) Cytotoxicity against JIMT1 target cells after 6 h (A) and 20 h (B) represents 4 repeats. (C) Representative experiment of cytotoxicity against BW target cells after 6 h represents 4 repeats. (D) Summary of BW cytotoxicity after 6 h represents 10 repeats. (E) Representative experiment of cytotoxicity against BW target cells after 20 h represents 6 repeats. (F) Summary of BW cytotoxicity at 20 h represents 10 repeats.

Across both tumor models, BE-conditioned EVs were associated with higher levels of target cell death compared to naive NK92-EVs. In the solid tumor model (JIMT1), this enhancement in cytotoxicity showed a concentration-dependent trend (Figure 4A, 4B). While untreated EVs induced modest killing, EVs derived from NK cells conditioned with 1 µM BE were associated with an increased cytotoxic effect. This phenotype was even more pronounced in the hematologic model (BW). In a 6-hour-long incubation (Figure 4C, 4D), EVs from the high-dose BE condition were associated with an increase in target cell death, consistent with enhanced early killing activity mediated by EVs. The strongest effects were observed after 20 hours (Figure 4E, 4F). Notably, we observed that 1 µM BE-EVs repeatedly achieved maximal cytotoxicity and outperformed the untreated and low-dose (1 pM) groups across all vesicle-to-cell ratios.

To determine whether BE exposure generally confers cytotoxic activity to EVs regardless of parental cell type, we next tested EVs derived from BE-conditioned PANC1 cells as a non-NK EV control. In contrast to BE-conditioned NK-derived EVs, PANC1-BE-EVs did not induce detectable JIMT1 killing at either 4 or 18 h, and no significant increase in target-cell death was observed across the tested EV inputs (Figure S3). These data indicate that BE exposure alone is not sufficient to generate cytotoxic EVs from a non-NK cellular source.

Collectively, these results are consistent with the conclusion that higher-intensity BE conditioning promotes the generation of EVs with increased anti-tumor activity in these assay systems.

**Figure 5.**
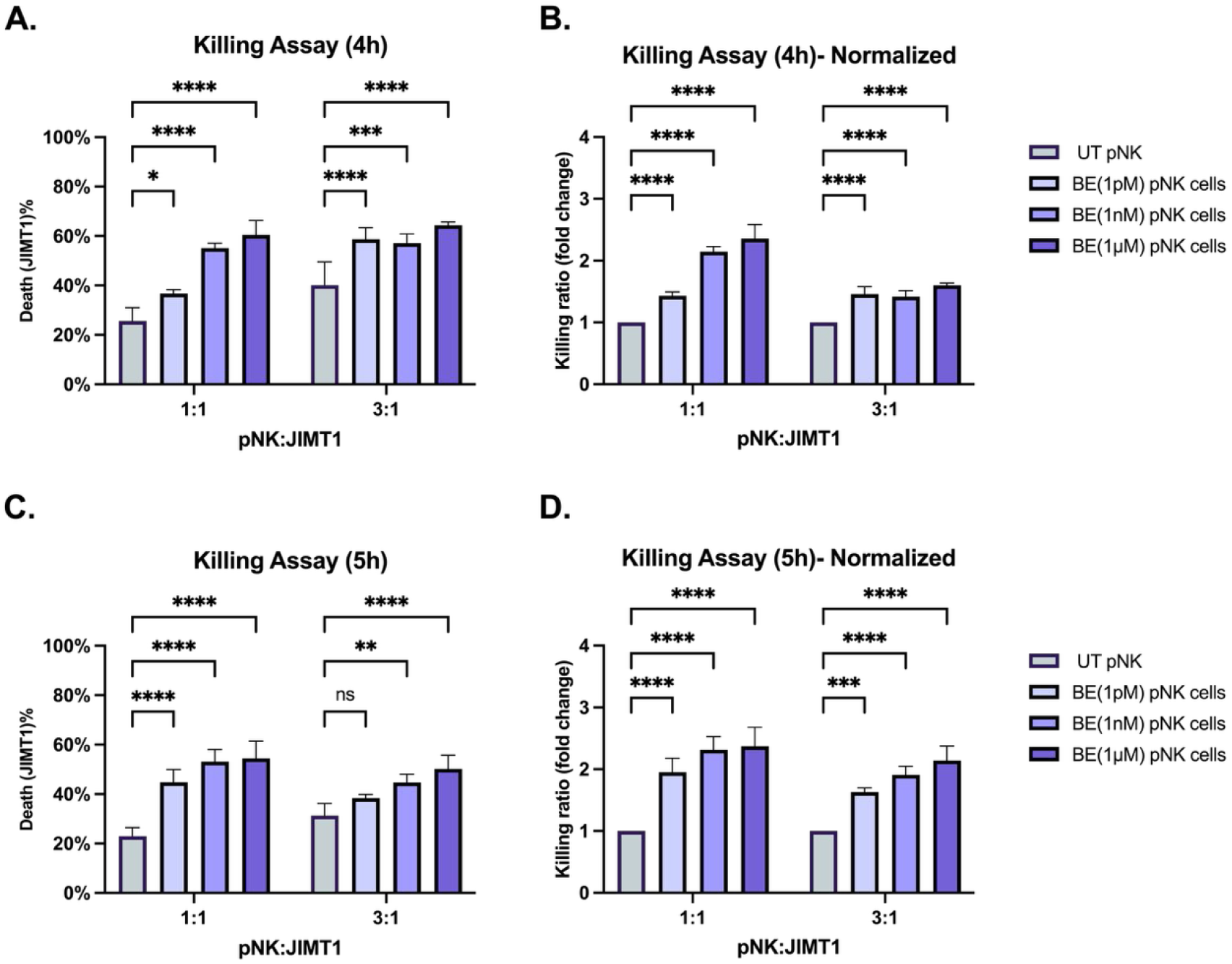
BE enhances primary NK cell-mediated killing of JIMT1 cells. Human primary NK (pNK) cells were co-cultured with JIMT1 target cells at pNK: JIMT1 ratios of 1:1 and 3:1 in the absence or presence of BE (1 pM, 1 nM, 1 µM). Target-cell death was assessed by flow cytometry as the percentage of DAPI⁺ JIMT1 cells. (A) Percentage of dead JIMT1 cells after 4 h of co-culture, represents 4 repeats. (B) Killing ratio at 4 h, normalized to the untreated pNK + JIMT1 control within each pNK: JIMT1 ratio. (C) Percentage of dead JIMT1 cells after 5 h of co-culture, represents 4 repeats. (D) Killing ratio at 5 h, normalized to the untreated pNK + JIMT1 control within each pNK: JIMT1 ratio. Data are presented as mean ± SD. Statistical significance was determined by two-way ANOVA followed by Dunnett’s multiple comparisons test comparing each BE-treated condition to the untreated control within the corresponding pNK: JIMT1 ratio.

### BE-mediated enhancement of NK cytotoxicity extends to primary human NK cells

To investigate whether the BE-associated increase in cytotoxicity observed in NK-92 cells could also be detected in primary human NK cells, short-term killing assays were performed using primary NK (pNK) cells co-cultured with JIMT1 breast cancer cells. pNK cells were incubated with JIMT1 cells at effector-to-target ratios of 1:1 and 3:1 in the absence or presence of BE (1 pM, 1 nM, 1 µM), and target-cell death was assessed by flow cytometry after 4 and 5 hours of co-culture.

At both time points, BE-treated pNK cells exhibited increased killing of JIMT1 cells compared with untreated controls. The effect was most pronounced at the 1:1 effector-to-target ratio, where baseline cytotoxicity was lower and BE treatment resulted in a clear increase in the proportion of dead JIMT1 cells. Consistent with this observation, normalization to the untreated pNK control revealed an approximately two-fold increase in killing activity across BE-treated conditions. At the 3:1 ratio, pNK cells displayed higher basal cytotoxicity, and BE treatment was associated with a more modest but reproducible increase in target-cell death.

Overall, the enhanced killing observed following BE exposure was evident at both 4 and 5 hours, indicating that BE can rapidly augment the cytotoxic activity of primary NK cells. These findings suggest that the BE-associated increase in tumor cell killing is not limited to the NK-92 cell line and may also be observed in primary human NK cells.

### BE-conditioned NK-EVs prime naïve NK-92 cells toward enhanced cytotoxic activity

Having shown that BE conditioning enhances the cytotoxic activity of NK-92-derived EVs, we next asked whether these vesicles could also act on naïve NK-92 cells and modulate their subsequent effector function. To separate EV-mediated priming of recipient NK cells from direct EV-mediated tumor killing, naïve NK-92 cells were incubated for 24 h with PBS, particle-normalized EVs from untreated NK-92 cells (NK92-EVs), or EVs from NK-92 cells conditioned with BE (1 pM, 1 nM, 1 µM); direct BE (1 µM) was included as a control for carry-over of free peptide. Cells were then washed extensively and co-cultured with JIMT1 target cells at NK-92:JIMT1 ratios of 1:1 and 3:1, and both tumor-cell death and NK-92 CD107a surface mobilization were assessed after 4 and 18 h.

Priming with EVs enhanced the subsequent cytotoxicity of naïve NK-92 cells against JIMT1 targets. At 4 h, all EV- and BE-primed conditions produced significantly greater target-cell death than PBS-primed controls at both effector-to-target ratios (Figure 6A), and this advantage over PBS was retained at 18 h, most consistently at the 1:1 ratio where baseline killing was lower (Figure 6C). Across the conditioning doses, the killing response trended upward with dose, with EVs from the 1 µM BE condition and the direct-BE control among the most active, while EVs from the 1 nM condition produced a comparable enhancement (Figure 6A, 6C).

**Figure 6.**
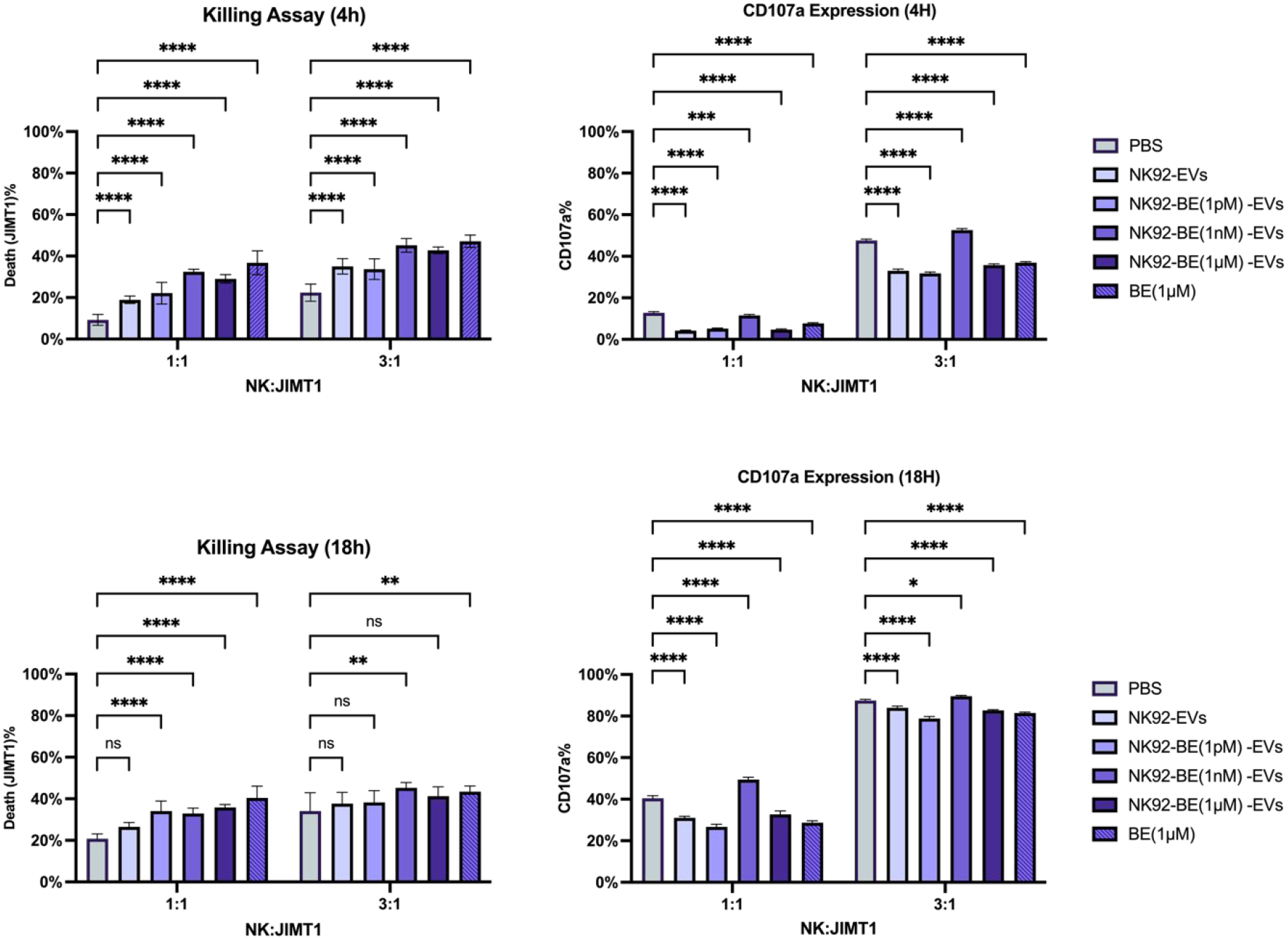
BE-conditioned NK-EVs prime naïve NK-92 cells and enhance secondary killing of JIMT1 cells. Naïve NK-92 cells were incubated for 24 h with PBS, NK92-EVs, or EVs from NK-92 cells conditioned with BE (1 pM, 1 nM, 1 µM), or with direct BE (1 µM), then washed and co-cultured with JIMT1 targets at NK-92:JIMT1 ratios of 1:1 and 3:1. **(A, C)** Percentage of dead (DAPI⁺) JIMT1 cells after 4 h (A) and 18 h (C) of co-culture. **(B, D)** Percentage of CD107a⁺ NK-92 cells after 4 h (B) and 18 h (D). Data are mean ± SD; n = 4. Statistical significance was determined by two-way ANOVA with selected multiple comparisons versus the PBS-primed control within each effector-to-target ratio (ns, not significant; *p < 0.05; ***p < 0.001; ****p < 0.0001).

Degranulation, in contrast, showed a distinct dose dependence. Whereas priming with NK92-EVs, 1 pM - or 1 µM-BE-EVs, and direct BE was generally associated with CD107a levels at or below those of PBS-primed cells, priming with 1 nM BE-EVs was uniquely associated with CD107a mobilization above the PBS control at 4 h (3:1) and at 18 h (both ratios) (Figure 6B, 6D). Thus the intermediate (1 nM) conditioning dose was the only condition in which enhanced killing was accompanied by increased surface CD107a.

Together, these findings indicate that BE-conditioned NK-EVs can act not only as direct cytotoxic effectors but also as intercellular mediators that raise the effector state of recipient naïve NK cells. The observation that enhanced killing was broadly transferable across conditioning doses, whereas increased degranulation was confined to the 1 nM-BE-EV condition, suggests that EV-mediated priming and classical granule exocytosis are separable: consistent with our earlier finding that direct BE enhances NK-92 killing without increasing CD107a (Figure 1E, 1F), most primed conditions augmented cytotoxicity through a degranulation-independent route, while an intermediate conditioning dose additionally licensed a classical degranulation response in recipient cells.

## Discussion

This study explores a neuroimmune mechanism by which β-endorphin (BE), a neuroendocrine mediator associated with positive physiological states, may modulate anti-tumor immunity by shaping NK cell function and secretory activity. BE exposure was associated with enhanced NK-92–mediated tumor killing, and importantly, this increased cytotoxicity persisted even in the absence of direct NK–target cell contact, indicating a prominent contribution of secreted effector components. The addition of primary human NK cell data further supports the biological relevance of this effect and suggests that BE-mediated enhancement of NK cytotoxicity is not restricted to the NK-92 cell-line model.

Mechanistically, the data suggest that BE does not primarily increase EV production but instead alters EV functional composition. While EV yield and size distribution remained stable across conditions, BE exposure was associated with enrichment of cytotoxic proteins, including granzyme B and perforin, within NK-derived EVs and with increased EV-mediated tumor cell killing. These findings support the idea that neuroendocrine signals can qualitatively reshape the cytotoxic secretome of NK cells.

Interestingly, enhanced tumor cell death occurred despite reduced CD107a mobilization, suggesting that BE may shift NK cytotoxic activity away from classical synapse-coupled degranulation toward alternative effector pathways. Consistent with this interpretation, conditioned media from BE-treated NK cells retained cytotoxic activity, while bulk IFN-γ levels remained largely unchanged, indicating that the observed effects are unlikely to be explained solely by soluble cytokine secretion.^32,33^

Taken together, these findings support a model in which BE can tune NK anti-tumor activity not only through direct modulation of effector cells but also by remodeling the vesicular arm of the NK secretome. In this context, NK-derived EVs may act as transferable cytotoxic units that mediate contact-independent tumor cell killing. The absence of comparable killing by PANC1-BE-EVs strengthens this interpretation by suggesting that the cytotoxic phenotype depends on the NK cellular origin and its pre-existing cytotoxic machinery.

This pattern supports a model in which BE signaling may influence the sorting and packaging of cytotoxic machinery into vesicles, potentially via altered trafficking through endosomal compartments, secretory lysosome pathways, or EV-loading mechanisms. Mechanistically, opioid receptor signaling on NK cells could plausibly modulate intracellular pathways that govern granule maturation, cytotoxic protein abundance, or cargo routing into multivesicular bodies and into released EVs. While the present dataset does not identify the molecular link, it narrows the likely mechanism toward cargo selection and compartmentalization rather than changes in vesicle output.

BE-EVs enhanced killing in both solid and hematologic cancer models, with a more pronounced phenotype in the hematologic setting. This may reflect differences in baseline susceptibility to vesicle-delivered cytotoxic proteins, EV uptake efficiency, membrane repair capacity, or intrinsic apoptosis thresholds between solid and hematologic tumor cells. Alternatively, it may indicate that BE conditioning changes not only EV cargo abundance (e.g., Granzyme B/Perforin) but also EV surface features that govern binding and internalization, which could vary by target cell type.

The primary NK cell results add an important validation layer to the NK-92 findings. In short-term co-culture assays, BE-treated primary human NK cells showed increased killing of JIMT1 cells, particularly under conditions in which basal cytotoxicity was lower. This suggests that BE can rapidly augment NK-mediated tumor killing in a primary human setting, although the magnitude and durability of this effect will need to be tested across additional donors and experimental conditions.

These findings support the concept that neuroendocrine signals can shape anti-tumor immunity at two complementary levels: by directly enhancing NK cell cytotoxic function and by remodeling the secreted effectors released by those cells. In our system, BE exposure was associated with stronger NK-–mediated killing across assay formats, indicating a robust increase in effector potency. In parallel, this functional shift extended into the cell-free compartment, where EV output remained broadly stable, but EV cargo and per-particle cytotoxic activity increased. Together, the data suggest that BE can “tune” anti-tumor activity both within the effector cell and through transferable vesicular units, providing a mechanistic foothold for psychoneuroimmunology by linking an endogenous peptide signal to measurable amplification of NK-driven cytotoxicity.

Several limitations should be considered when interpreting these results. First, although the inclusion of primary NK cells supports the relevance of the NK-92 observations, most mechanistic EV experiments were performed using NK-92 cells, which are clinically relevant but may not fully recapitulate primary NK cell biology. Therefore, future experiments should evaluate whether BE-conditioned primary NK cells generate EVs with similar cytotoxic cargo enrichment and anti-tumor activity. Furthermore, profiling EV surface proteins and testing uptake kinetics across different targets will be necessary to interpret the differential susceptibility observed between solid and hematologic tumor models and to strengthen mechanistic claims.

The mechanistic basis of BE signaling also needs further investigation, demonstrating receptor involvement (e.g., opioid receptor blockade) and mapping downstream signaling changes. To address this, exploring intracellular signaling pathways that may directly link opioid receptor activation to EV cargo selection and routing will be required. Finally, to advance the translational potential of these findings, BE-conditioned EVs should be evaluated across a broader panel of tumor models and in suitable in vivo systems. Establishing this “neuroendocrine priming” effect in primary cells and primary NK-derived EVs is a critical next step toward developing it as a viable *ex vivo* conditioning strategy to enhance EV potency prior to therapeutic use.

In summary, our data support a model in which BE exposure is associated with enhanced NK anti-tumor activity and a parallel shift toward contact-independent cytotoxic effector function. BE treatment increased NK-mediated killing across time points, despite reduced CD107a mobilization, suggesting that the functional enhancement is not fully captured by classical degranulation readouts and may involve alternative cytotoxic pathways.

Beyond the mechanistic findings, these results offer a conceptual framework for how positive-state-linked neuroendocrine signals could shape anti-tumor immunity before immune cells physically encounter transformed targets.

Together, these data are consistent with the possibility that BE can bias NK cells toward a contact-independent effector program, in which cytotoxic potential is exported into vesicular units that can act at a distance from the parent cell. Although broader validation in primary NK cells, primary NK-derived EVs and suitable *in vivo* models are necessary for extrapolation to physiological aspects of cancer prevention, these findings offer a mechanistic basis for psychoneuroimmunology. They suggest that neuroendocrine signals associated with positive physiological states may “precondition” NK cells to enhance EV-mediated cytotoxicity, thereby increasing early immune responses against developing malignant cells.

The EV-priming experiment adds an additional layer to this model by showing that NK-derived EVs may influence not only tumor cells but also other NK cells. Following removal of the priming stimulus, NK-92 cells exposed to NK-derived EVs displayed enhanced secondary killing of JIMT1 targets, suggesting that EVs can transfer a functional effector state to recipient NK cells. This supports the possibility that NK-EVs participate in immune-to-immune communication and may amplify anti-tumor activity beyond the original BE-exposed NK cell population.

Notably, EV-mediated enhancement of killing was not uniformly accompanied by increased CD107a mobilization. Most priming conditions enhanced cytotoxicity without increasing CD107a, whereas 1 nM BE-EVs uniquely increased CD107a expression. This finding strengthens the conclusion that increased NK cytotoxicity in this system is not explained solely by classical degranulation. Instead, BE-conditioned EVs may activate separable cytotoxic programs in recipient NK cells, with some conditions enhancing killing through degranulation-independent mechanisms and the intermediate 1 nM BE-EV condition additionally promoting a classical degranulation-associated response.

In summary, our data support a model in which BE enhances NK anti-tumor activity through multiple connected mechanisms: direct enhancement of NK-mediated killing, remodeling of NK-EV cytotoxic cargo, increased EV-mediated tumor killing, and secondary priming of naïve NK cells. Together, these findings suggest that BE does not simply increase general NK activation, but may bias NK cells toward a secretory and vesicular effector program capable of acting both on tumor targets and on neighboring immune cells. Although broader validation in primary NK cells, primary NK-derived EVs, and suitable in vivo models will be required, these results provide a mechanistic framework for how positive-state-linked neuroendocrine signals may precondition NK cells and their EVs to amplify anti-tumor immunity.

